# <u>M</u>inimum <u>I</u>nformation for <u>Re</u>usable <u>A</u>rthropod <u>A</u>bundance <u>D</u>ata (MIReAAD)

**DOI:** 10.1101/429142

**Authors:** Samuel Rund, Kyle Braak, Lauren Cator, Kyle Copas, Scott J. Emrich, Gloria I. Giraldo-Calderón, Michael A. Johansson, Naveed Heydari, Donald Hobern, Sarah A. Kelly, Daniel Lawson, Cynthia Lord, Robert M MacCallum, Dominique G. Roche, Sadie J. Ryan, Dmitry Schigel, Kurt Vandegrift, Matthew Watts, Jennifer M. Zaspel, Samraat Pawar

## Abstract

Arthropods play a dominant role in natural and human-modified terrestrial ecosystem dynamics. Spatially-explicit population time-series are crucial for statistical or mathematical models of these dynamics and assessment of their veterinary, medical, agricultural, and ecological impacts. Arthropod data have been collected world-wide for over a century, but remain scattered and largely inaccessible. With the ever-present and growing threat of arthropod vectors of infectious diseases and pest species, there are enormous amounts of historical and ongoing surveillance. These data are currently reported in a wide variety of formats, typically lacking sufficient metadata to make reuse and re-analysis possible. We present the first minimum information standard for arthropod abundance. Developed with broad stakeholder collaboration, it balances sufficiency for reuse with the practicality of preparing the data for submission. It is designed to optimize data (re-)usability from the “FAIR,” (Findable, Accessible, Interoperable, and Reusable) principles of public data archiving (PDA). This standard will facilitate data unification across research initiatives and communities dedicated to surveillance for detection and control of vector-borne diseases and pests.

## Introduction

Arthropods play a dominant role in the dynamics of practically all natural and human-modified terrestrial ecosystems^1–3^, and have significant economic and health effects. For example, certain insects provide significant economic benefits (*e.g.* pollination) exceeding $57 billion a year to the United States alone^4^. Meanwhile, invasive insects cost an estimated $70 billion dollars per year globally^5^ and insect pests may reduce agricultural harvests by up to 16%, with an equal amount of further losses of harvested goods^6^. Particularly noteworthy is a subset of arthropods that are disease vectors, transmitting pathogens to and between animals as well as plants. Vector-borne diseases cause billions of dollars in crop and livestock losses, every year^7–9^. In humans, vector borne diseases account for more than 17% of all infectious diseases (*e.g.* malaria, Chagas, dengue, and leishmaniasis, Zika, West Nile, Lyme disease, and sleeping sickness), with hundreds of thousands of deaths, hundreds of millions of cases, and billions of people at risk, annually^10,11^.

The current economic and health burden of arthropod pests, exacerbated by invasive species, and uncertain effects of climate change^12,13^, has driven significant research programs and data collection efforts. These include crop pest, mosquito, and tick survey and reporting initiatives^14–18^, citizen science projects^19–21^, and digitization of museum specimen data^22,23^, all yielding a rich and growing trove of field-based data spanning multiple spatial and temporal scales. Monitoring arthropod abundance (*e.g.* Figure 1) in different disciplines (*e.g*., biodiversity research, pest-control assessment, vector-borne disease monitoring, or pollination research) uses similar techniques, with similar objectives: to quantify abundance, phenology and geographical ranges of target arthropod species. Despite a growing number of data collections, they are often not reusable, or comparable to similar data, due to a lack of standardization and metadata. In contrast, the advent of the deposition of data from high-throughput technologies (*e.g.* NCBI and GenBank), data and code sharing, and other practices to improve transparency and reusability of research results are increasing rapidly across the sciences^24–29^. Furthering these advances through standardization and public archiving of arthropod abundance data can bring significant benefits, including (1) supporting empirical parameterization and validation of mathematical models (*e.g.* of pest or disease emergence and spread), (2) validation of model predictions, (3) reduction in the duplication of expensive empirical research, and (4) revealing new patterns and questions through meta-analyses^30–33^. This will also lead to substantial public benefit through improved human, animal, plant, and ecosystem health, and reduced economic costs.

**Figure 1.**
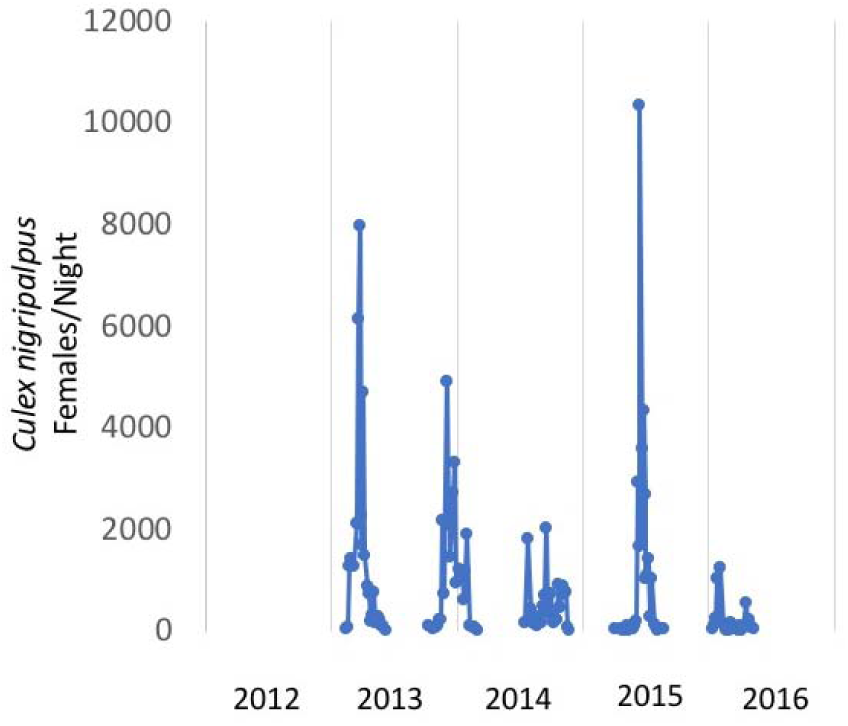
Example population abundance time-series.

A key impediment to the re-use of these data is the lack of adequate metadata or data descriptors (*i.e.* data about the data)^34–37^. In general, for data to be most valuable to the scientific community, they should meet the FAIR Principles – they should be Findable, Accessible, Interoperable and Reusable – and delineate the key components of good data management and stewardship practices^38,39^. Data are Findable and Accessible when they are archived and freely downloadable from an online public data repository that is indexed and easily searchable. Interoperability and reusability describe the ease with which humans or computer programs can understand the data (*e.g.* via metadata) and explore/re-use them across a variety of non-proprietary platforms. Even when data are available, metadata for arthropod abundance data are often absent or not readily interpretable, limiting their reusability at a fundamental level.

## Results

### A minimum information standard for arthropod abundance data

Here, we present a Minimum Information for Reusable Arthropod Abundance Data (MIReAAD) standard for reporting primarily longitudinal (repeated, temporally explicit) field-based collections of arthropods. In the same manner as has been developed in other biological disciplines^40–45^, this standard is “minimum” because it defines the necessary minimal information required to understand and reuse a dataset without consulting any further text, materials, or methods^46^. MIReAAD is designed to facilitate data archiving efforts of publishers and field researchers. It is not a data model and therefore does not define controlled vocabularies, or specific field titles, but should be easy to understand, and interpret by the wider scientific community^46^.

The minimal standards are separated into two components, metadata and data. For each component, we provide a description of the information that should be included, recommendations for how to make that information as useful as possible, and examples. The metadata component (Table 1) includes information for the origin of the data set (*e.g*. study information and licensing for usage). The second component (Table 2) lists and describes specific data fields that should be included in data collection sheets. We also provide recommendations and examples to demonstrate how these recommendations can be implemented. MIReAAD was designed to match the data that are generally collected by academic researchers and surveillance initiatives, and can serve as a checklist for important information that needs to be recorded but is often unintentionally omitted (*e.g.* Figure 2A). By adhering to MIReAAD standards, omissions and ambiguity can be avoided even if the data are shared in different formats (Figure 2B and C). Finally, we identify common problems likely to be encountered across all the MIReAAD metadata and data fields, and data quality standards that can be employed to avoid confusion (Box 1).

**Table 1.**
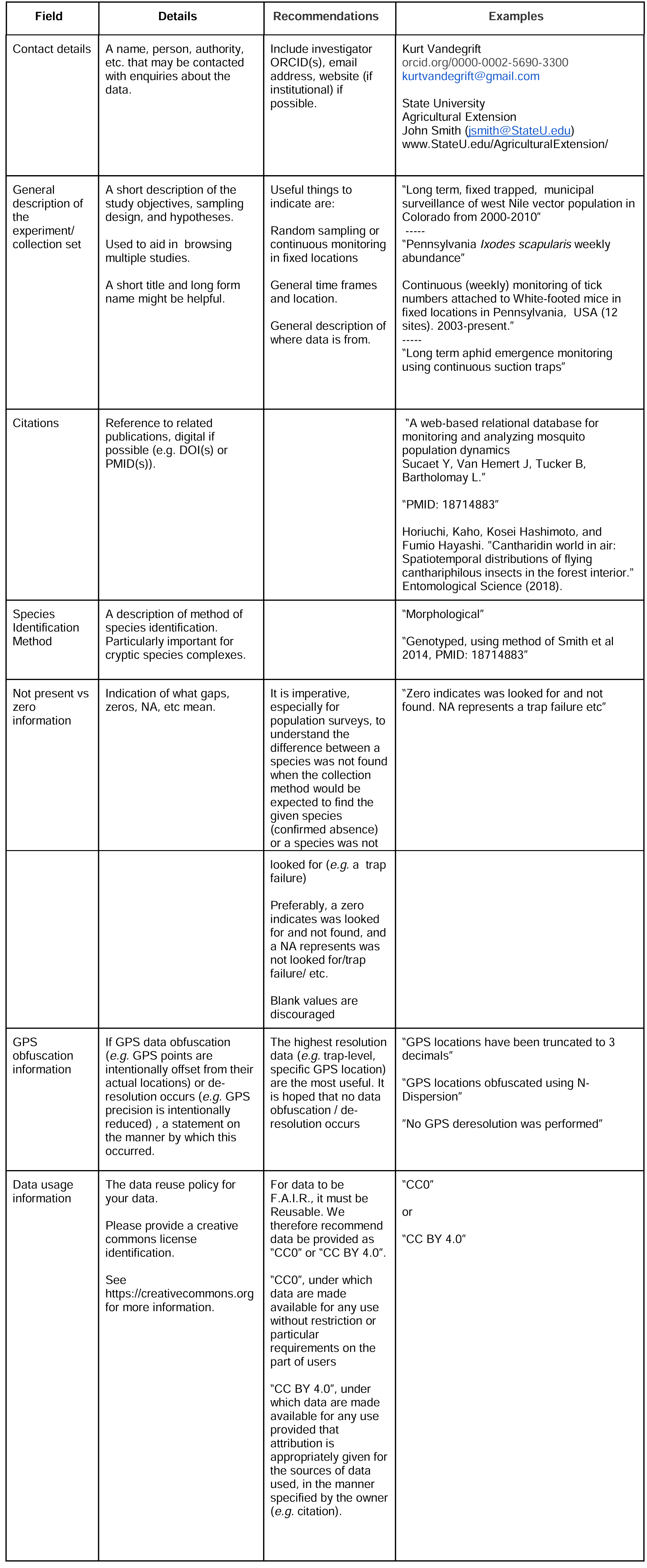
The MIReAAD Study Information (Resource metadata) fields. The information in this table should be included with every data submission, for example by including data in the file header as demonstrated in Data Files 1-4.

**Table 2.**
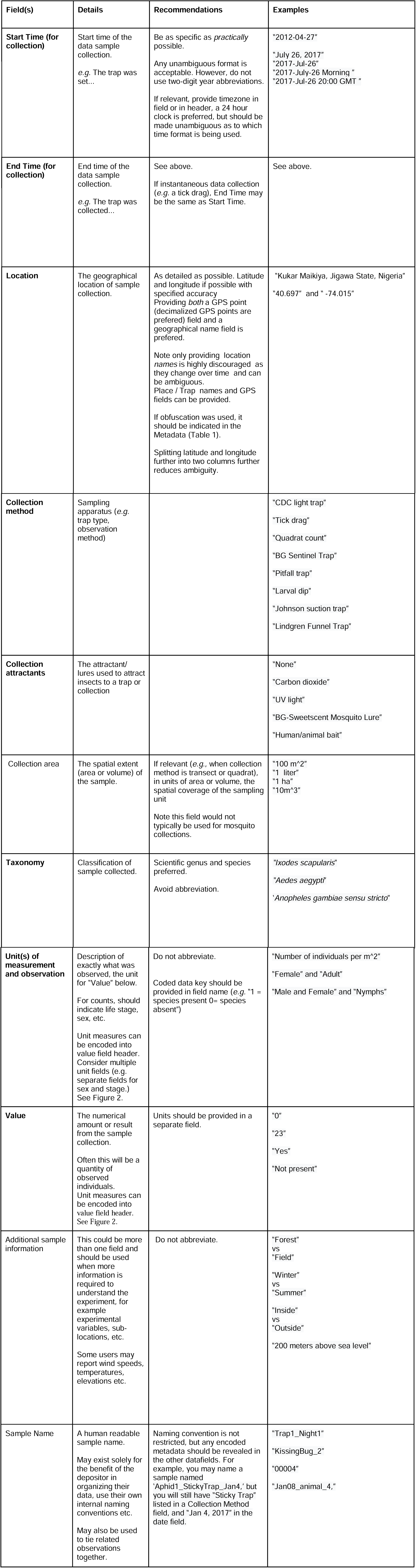
The MIReAAD data fields. Fig 1B provides an annotated example.

| Field(s) | Details | Recommendations | Examples |
| --- | --- | --- | --- |
| **Start Time (for collection)** | Start time of the data sample collection.<br>e.g. The trap was set... | Be as specific as *practically* possible.<br>Any unambiguous format is acceptable. However, do not use two-digit year abbreviations.<br>If relevant, provide timezone in field or in header, a 24 hour clock is preferred, but should be made unambiguous as to which time format is being used. | "2012-04-27"<br>"July 26, 2017"<br>"2017-Jul-26"<br>"2017-July-26 Morning "<br>"2017-Jul-26 20:00 GMT " |
| **End Time (for collection)** | End time of the data sample collection.<br>e.g. The trap was collected... | See above.<br>If instantaneous data collection (e.g. a tick drag), End Time may be the same as Start Time. | See above. |
| **Location** | The geographical location of sample collection. | As detailed as possible. Latitude and longitude if possible with specified accuracy<br>Providing *both* a GPS point (decimalized GPS points are preferred) field and a geographical name field is preferred.<br>Note only providing location *names* is highly discouraged as they change over time and can be ambiguous.<br>Place / Trap names and GPS fields can be provided.<br>If obfuscation was used, it should be indicated in the Metadata (Table 1).<br>Splitting latitude and longitude further into two columns further reduces ambiguity. | "Kukar Maikiya, Jigawa State, Nigeria"<br>"40.697" and "-74.015" |
| **Collection method** | Sampling apparatus (e.g. trap type, observation method) |  | "CDC light trap"<br>"Tick drag"<br>"Quadrat count"<br>"BG Sentinel Trap"<br>"Pitfall trap"<br>"Larval dip"<br>"Johnson suction trap"<br>"Lindgren Funnel Trap" |
| **Collection attractants** | The attractant/ lures used to attract insects to a trap or collection |  | "None"<br>"Carbon dioxide"<br>"UV light"<br>"BG-Sweetscent Mosquito Lure"<br>"Human/animal bait" |
| Collection area | The spatial extent (area or volume) of the sample. | If relevant (e.g., when collection method is transect or quadrat), in units of area or volume, the spatial coverage of the sampling unit<br>Note this field would not typically be used for mosquito collections. | "100 m<sup>2</sup>"<br>"1 liter"<br>"1 ha"<br>"10m<sup>3</sup>" |
| **Taxonomy** | Classification of sample collected. | Scientific genus and species preferred.<br>Avoid abbreviation. | "*Ixodes scapularis*"<br>"*Aedes aegypti*"<br>'*Anopheles gambiae sensu stricto*' |
| **Unit(s) of measurement and observation** | Description of exactly what was observed, the unit for "Value" below.<br>For counts, should indicate life stage, sex, etc.<br>Unit measures can be encoded into value field header. Consider multiple unit fields (e.g. separate fields for sex and stage.) See Figure 2. | Do not abbreviate.<br>Coded data key should be provided in field name (e.g. "1 = species present 0= species absent") | "Number of individuals per m<sup>2</sup>"<br>"Female" and "Adult"<br>"Male and Female" and "Nymphs" |
| **Value** | The numerical amount or result from the sample collection.<br>Often this will be a quantity of observed individuals. Unit measures can be encoded into value field header. See Figure 2. | Units should be provided in a separate field. | "0"<br>"23"<br>"Yes"<br>"Not present" |
| Additional sample information | This could be more than one field and should be used when more information is required to understand the experiment, for example experimental variables, sub-locations, etc.<br>Some users may report wind speeds, temperatures, elevations etc. | Do not abbreviate. | "Forest"<br>vs<br>"Field"<br>"Winter"<br>vs<br>"Summer"<br>"Inside"<br>vs<br>"Outside"<br>"200 meters above sea level" |
| Sample Name | A human readable sample name.<br>May exist solely for the benefit of the depositor in organizing their data, use their own internal naming conventions etc.<br>May also be used to tie related observations together. | Naming convention is not restricted, but any encoded metadata should be revealed in the other datafields. For example, you may name a sample named 'Aphid1_StickyTrap_Jan4,' but you will still have "Sticky Trap" listed in a Collection Method field, and "Jan 4, 2017" in the date field. | "Trap1_Night1"<br>"KissingBug_2"<br>"00004"<br>"Jan08_animal_4," |

**Figure 2.**
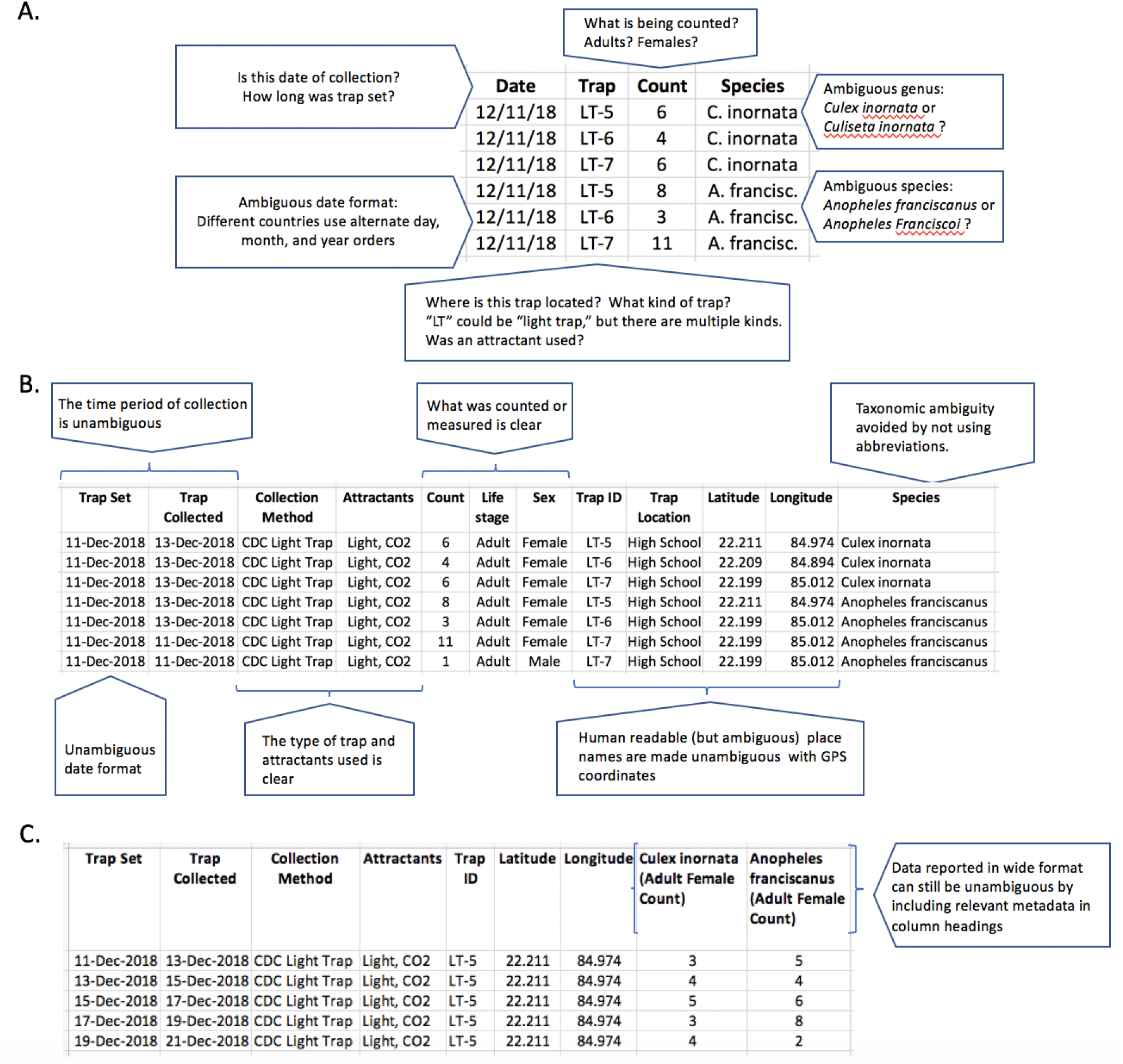
MIReAAD reduces data ambiguity. A. Seemingly clean data can still lack key information or have ambiguous metadata, hindering data reuse. B. MIReAAD compliant data includes the metadata necessary for data reuse and removes ambiguity. C. Note data can be formatted differently, but still be MIReAAD complaint, such as by presenting data in a wide format

##### Box 1. Data quality standards

###### No abbreviations

Abbreviations (including in columns names) are ambiguous, with the exception of measurement units (*e.g.* centigrade and meters).

###### No external legend/key files

While repetitive, all data should be explicitly given within the data table. Separate files mapping ID numbers to GPS locations, full species names, etc., should be avoided. In addition, rich metadata is essential for good data discovery and reuse.

###### Unambiguous dates

Because of country-level differences in date formats, data should be reported with 4 digit years, and months provided alphabetically and not numerically (*e.g.* 4-Jun-2017 or Nov 12, 2015).

###### Machine-readable file formats

Data should be provided in non-proprietary machine readable formats such as comma-separated text files. PDFs and multiple spreadsheets in the same document should be avoided.\

###### No font styling or subsection headings

Formatting (color, bold, italics, subscripts, sheet tab names, *etc.*) should not be required for understanding the data. Subsection headings should not be required to understand data; every line of data should be interpretable in isolation from any other line of data.

###### Highest precision possible

Data should be provided at the highest temporal, spatial, numerical, and taxonomic resolution available. If location (*e.g.*, geographical coordinate) data need to be presented at a lower resolution than available for privacy reasons, this should be made clear in the submission in Study Information (Resource Metadata; Table 1).

###### Language

Once data are ready to be deposited/submitted, all fields and data are preferably written in English. This will allow researchers and data curators worldwide to understand and reuse the data. Use of other languages is better than not publishing data. Please avoid introducing data reuse barriers through incomplete translation. For example, non-English field names in an English-language submission.

### Examples

Below we provide three examples to illustrate MIReAAD compliant data (linked to Supplemental Data Files 1-4, respectively). Researchers can use these data sheets as a basis for formatting their own data. In these examples, note that all data meet the data quality standards of Box 1; are adequately described, have columns labeled, *etc*. to eliminate ambiguity (even if the data appear repetitive; for example, the sex and life stage are repeated in every row). Examples 1 and 2 should be sufficient for most data generators. Example 3 (Data Files 3-4) demonstrates a more complex data collection scenario.

#### 1. Long-format trapping data

Each row captures count data for a single species’ occurrence in a given sampling event. This illustrates an example of the most common mosquito collection protocol. [<u>Sup Datasheet 1</u>]. Also see Figure 2B.

#### 2. Wide format trapping data

Each row captures count data from a given sampling event. Each identified taxonomic group is identified in a separate column. An ‘additional sample information’ field, ‘sub-location,’ has been added to describe the various locations around the village where collections were made. [<u>Sup Datasheet 2</u>]. This illustrates an example of adult mosquito populations that have been tracked over time and in specific locations. Also see Figure 2C.

#### 3. Complex trapping data scenario

Tick surveillance performed using tick drags and flags and collections of ectoparasites on trapped mice. The tick drags/flags report three life stages independently (adult, larvae, and nymph) [<u>Sup Datasheet 3</u>]. Larvae are only identified to the genus, while adults and nymphs are identified to the species. Observations of different life stages and sexes are preferably documented in separate records. A Sample Name is used to help link these records (but would not be necessary.) The mouse survey uses an additional sample information field to record the sex of the trapped mouse from which the parasites were collected [<u>Sup Datasheet 4</u>].

## Discussion

### MIReAAD as the path to FAIR data principles

We designed MIReAAD to achieve a balance between standards that are too onerous for data generators and standards that are sufficient to ensure at least minimal reusability^31,40^. Like all minimum standards, MIReAAD only aims at ensuring data ‘Reusability’. However, ultimately this will promote the implementation of data models — the explicit definition of data field names, data formats (*e.g.*, for dates and GPS locations), and controlled vocabularies (*e.g.*, the Darwin Core^47^). Data models enable ‘Interoperability’, and in turn facilitate structured databases, public repositories, and development of data analysis tools^46,48^. Deposition in open databases make data ‘Findable’ and ‘Accessible’^49–51^. MIReAAD compliant data contain sufficient information for established aggregators/databases such as VectorBase and SCAN (Symbiota Collections of Arthropods Network^52^) to process and store the data in a standardized data model [*e.g.*, Darwin Core, a widely used universal data standard that supports opportunistic observation and collection data (occurrence core) as well as presence/absence and abundance data collected using strict and documented methodology (event core)^47^], and ultimately facilitate data transfer to even more comprehensive biodiversity databases [*e.g.* GBIF, which contains over one billion species occurrence records, from thousands of environmental, ecological, and natural resource investigations, including research on Arthropoda in numerous ecological and monitoring projects, allowing for study of changes and trends in populations.^51^]. Indeed, in Supplemental File 5, we provide an example of the mapping of data fields from this minimum information standard, to DarwinCore and GBIF. In this way, MIReAAD opens the door to FAIR data and more sophisticated methods to integrate data across many scales.

### Benefits to field researchers

It is essential that the benefits of a minimal data standard extend not just to data re-users, but also to the researchers who collect and generate data in the first place. MIReAAD provides a framework for data preparation that can help scientists achieve recognized professional merit for sharing data such as increased citation rates, academic recognition, opportunities for co-authorship, and new collaborations [sensu Roche et al. 2014^31^]. Large, deposited data sets can now themselves be standalone, citable “data papers” (*e.g. ^53–55^*) or even depositions without any traditional manuscript (but as an authored ‘digital product,’ with persistent identifiers, such as a DOI number), if desired. Data sets are increasingly recognized as valuable research outputs that count towards academic recognition and professional advancement (*e.g.* grants, interviews, and tenure). For example, several funders (*e.g.* United States National Science Foundation and Swiss National Science Foundation) have adopted or are in the process of adopting the Declaration on Research Assessments (DORA)^56^, offering further opportunities for data generators to gain recognition and publication credit for their work^57^. Also, an increasing number of funders are mandating public data access, and detailed data management plans are often required even at the grant proposal stage. Therefore, reporting data according to MIReAAD will provide a foundational pipeline for stipulating archival formats.

Furthermore, many data generators are also data users. Developing analyses that rely on standardized fields can facilitate the development of generalized analytical tools that can be easily extended to datasets beyond those that were collected by a single individual or lab. In this way, they can enable extensions of work that would otherwise not happen, such as comparisons of population dynamics in different locations or assessments of interspecies interactions. Adopting MIReAAD therefore can both help data generators reap the benefits of sharing data they have collected and enable them to more readily leverage data collected by others.

### Further MIReAAD applications and extensions

The creation of minimum information standards for these types of databases facilitates analyses of data at the scales that cannot be attained by a single individual or lab group. Linking records to additional information also extends the utility of these data to address population level questions. For example, a well-populated database presents opportunities to investigate interactions between populations of different species of arthropod that overlap in geography, but may be of interest individually to different realms of research. As a case in point, in the northeastern USA, *Agrilus plannipennis,* the Emerald Ash Borer (EAB), is a highly destructive invasive insect, monitored closely by both state and federal agencies for management^58^. Interestingly, EAB are creating lots of new habitat for carpenter bees, a species interaction that can be tracked and anticipated using large scale arthropod data.

Another example of the utility of linked data is for disease vectors. Data on insecticide resistance linked with time and place would be valuable for coordinating control strategies within and between nations and communities. Presence/absence data on infection levels would be helpful for tracking and investigating disease outbreaks, and dynamics. Standardization of these data would be particularly useful for pathogens that infect multiple vectors and hosts and would facilitate a “One Health” approach. Other important vector phenotypes that contribute to control and transmission such as pathogen susceptibility, biting preferences, and breeding behaviours could be measured over time and space.

We note that MIRreAAD is applicable not only to abundance measurements, but could be easily extended to any other kind of routinely sampled time-series field data. For example, in addition to aphid abundance, plant pathogen (such as mosaic virus) infection and insecticide resistance statuses of the aphids could be reported in MIRreAAD format.

## Conclusion

We present MIReAAD, a minimum information standard for representing arthropod abundance data. MIReAAD will facilitate collation and analyses of data at scales that cannot be attained by a single individual or lab, to address key questions across temporal and spatial scales, such as within and across-year phenology of abundance of target arthropod taxa over large geographical areas. This is particularly important given the pressing need to understand and predict the population dynamics of harmful (e.g., disease vectors and pests) as well as beneficial (e.g., pollinators, bio-control agents) arthropods in natural and human modified landscapes. This is the first step for achieving the broad benefits of FAIR data for arthropod abundance. We call on data generators, authors, reviewers, editors, journals, research infrastructures (*e.g.* data repositories) and funders to embrace MIReAAD as a standard to facilitate FAIR data use and compliance for arthropod abundance data.

## Author contributions

The project was conceptualized by Lauren Cator and Samraat Pawar. The original draft was prepared by Michael A. Johansson, Samuel S.C. Rund, Naveed Heydari, Kurt Vandegrift, Matthew Watts, and Samraat Pawar. Visualization was prepared by Kurt Vandegrift, Samuel S.C. Rund, Samraat Pawar, and Michael A. Johansson. Review & Editing was performed by all the authors.

## Acknowledgements

The seeds of this effort were planted in 2016 at a meeting of VectorBiTE, which is a cross-disciplinary research coordination network (RCN) for disease vectors. Samuel S.C. Rund, Matthew Watts, Kurt Vandegrift, Naveed Heydari, Cynthia Lord, Michael Johansson, Samraat Pawar, and Sadie J. Ryan, received travel funding from NIH grant 1R01AI122284-01 and BBSRC grant BB/N013573/1 as part of the joint [NIH-NSF-USDA-BBSRC] Ecology and Evolution of Infectious Diseases program.

Samuel S.C. Rund was funded by the Royal Society (NF140517). Rund, Daniel Lawson, Robert M. MacCallum, Sarah A. Kelly, Gloria I. Giraldo-Calderon and Scott J. Emrich were supported by the National Institute of Allergy and Infectious Diseases, National Institutes of Health, Department of Health and Human Services, under Contract No. HHSN272201400029C (VectorBase Bioinformatics Resource Center).

Kurt Vandegrift was funded by the National Science Foundation Ecology and Evolution of Infectious Diseases program (1619072).

Naveed Heydari and Sadie J. Ryan were funded by National Science Foundation (NSF DEB EEID 1518681).

Sadie J. Ryan was additionally funded by NIH 1R01AI136035-01, and CDC grant 1U01CK000510-01: Southeastern Regional Center of Excellence in Vector-Borne Diseases: the Gateway Program. This publication was supported by the Cooperative Agreement Number above from the Centers for Disease Control and Prevention. Its contents are solely the responsibility of the authors and do not necessarily represent the official views of the Centers for Disease Control and Prevention.

Jennifer M. Zaspel was funded by the National Science Foundation Division of Biological Infrastructure (NSF 1561448, NSF 1601957).

